# Differential colonization and succession dynamics of marine bacteria on different plastic polymers

**DOI:** 10.1101/2024.02.21.581331

**Authors:** Keren Davidov, Sheli Itzahri, Liat Anabel Sinberger, Matan Oren

**Affiliations:** Department of Molecular Biology, Ariel University, Ariel, Israel

**Author notes:** Equal authorship (Keren Davidov, Sheli Itzahri).

## Abstract

During the past decades since plastic was introduced to the world, marine microorganisms have been adapted for life on marine plastic debris, forming unique plastic-attached microbial communities. To date, little is known about the colonization and succession processes that take place on plastic surfaces in marine environments and how the plastic polymer type affects the plastic-attached microbiome composition. To address this knowledge gap, we examined the colonization and succession dynamics of marine bacteria on four common plastic polymers - PE, PP, PS, and PET-compared to glass and wood in a controlled seawater system under different temperatures. Using a simple experimental design, coupled with a long-read 16S rRNA metabarcoding pipeline and a set of complementary data analyses, we characterized the temporal trends in the composition of the bacterial microbiome developed on different surfaces over 2 - 90 days. By applying weighted gene co-expression network (WGCNA) analysis, we established co-occurrence networks and identified genera with specific succession signatures, significant enrichment on specific plastic polymers and/or strong intra-genus connections. Among them, members of genus *Alcanivorax* were significantly enriched on either PE or PP plastic surfaces as early as 2 days post-inoculation. *Alcanivorax* colonization preference to polyolefins was confirmed in colonization assays with pure *Alcanivorax* strains. Our research approach presented here may contribute to understanding how plastisphere communities are being formed and help identify taxa with specific adaptations to plastic surfaces.

## Introduction

Marine plastic pollution is a primary ecological concern. Like global climate change, it is noticeable in its effect on the biosphere and the life of many species. During the past seven decades since plastic was introduced to the marine environment, microorganisms have been adapted to live and thrive on marine plastic debris. Within minutes after arrival to the marine environment, plastic surfaces are colonized with marine bacteria [1], leading to the formation of an initial biofilm, which in turn develops into a rich community consisting of a variety of phototrophs and heterotrophs, predators, decomposers and symbionts [2]. Due to their unique taxonomic composition compared to the surrounding water, marine plastic debris and their biological cover are considered a separate ecosystem referred to as the “plastisphere” [3]. The microbial communities of the plastisphere are affected by different factors, including geography and season [4-7], immersion/incubation time [8, 9], salinity [10] and plastic weathering state [11, 12]. While the composition of the plastic-attached microbiome differs from that of natural organic debris [13], the effect of the plastic polymer type remains elusive and often undetectable [8, 14-16]. A few studies showed polymer-based variations in the early marine plastic microbiome, a few days to several weeks old [17-22]. However, statistically valid colonization and succession signatures of individual species within a naturally developed biofilm are missing.

In the present study, we examined the colonization and succession dynamics of marine bacteria on four common plastic polymers (Polyethylene - PE, Polypropylene - PP, polystyrene - PS and Polyethylene Terephthalate - PET) as well as on non-plastic surfaces (glass and wood) in a controlled seawater system. Using a simple experimental design, coupled with MinION nanopore-based, long-read 16S rRNA metabarcoding pipeline for bacteria and a set of complementary data analyses, we characterized the trends in the microbiome biodiversity and composition of the different surfaces. By applying weighted gene co-expression network (WGCNA) analysis on the relative abundance datasets of the samples, we have identified bacterial genera with relative surface- and time-specific colonization preferences.

## Methods

### Experiments setup

The experiments were carried out in a 70-litter semi-sterile seawater aquarium. Water temperature was regulated using a chiller and a heater. The first (will be referred to as the ‘main’) experiment was carried out at 28 ± 2°C. We performed another experiment to examine the effect of temperature on colonization dynamics. The second experiment was carried out at a higher temperature of 35 ± 3 °C. Water salinity was kept at ∼40 PPT in both experiments. The aquarium included two compartments divided by a semipermeable barrier, allowing free water flow but preventing the movement of large items from one side to the other (Supplementary Fig. S1). For the inoculation of plastic-associated marine bacteria, different plastic debris was collected from Herzliya Marina in the Mediterranean Sea (32° 09′ 38.8” N 34° 47′ 35.0” E). All items were washed with bacteria-free filtered artificial seawater (FASW). The items were then submerged in the aquarium compartment that received a constant inflow of sump-filtered sea water (Supplementary Fig. S1, right). Six days after the marine plastic trash items were introduced, new surfaces were added to the left compartment of the aquarium from which water is flowing back to the sump. The new surfaces included organza mesh bags (6 × 8 cm in size), each containing 7 gr of beads (3-5 mm in diameter) of either pristine (pre-processed, pure) PE, PP, PS, or PET plastic polymers, glass or wood. The second experiment, which was conducted at a higher temperature, included only PE, PET, glass, and wood. The begs (including replicates for each material according to the number of sampling time points) were hung in mid-aquarium water by thin fishing lines tied to the aquarium top cover (Supplementary Fig. S1, left).

### Sampling

Pieces from four plastic debris (D1-D4 in the main experiment and D5-D8 in the second experiment) were sampled at the beginning of the experiment. Sampling of the newly introduced beads was performed at 2, 7, 14, 30, and 90 days from the beginning of the main experiment and 7, 14, and 30 days from the beginning of the second experiment. At each time point, one organza bag of each tested material and half a litter of the surrounding aquarium water were sampled. Beads were carefully washed in FASW before processing. The water was filtered through a 0.22 µm polyethersulfone membrane using a 20 L/min pump (MRC). The beads and the membranes were subjected to DNA extraction as described below.

### DNA extraction

Pooled beads of each tested substrate, pieces of plastic debris, and membranes with water filtrate were subjected to DNA extraction using the phenol-chloroform method. The extraction was performed in 15 ml tubes containing ∼ 0.4 gr of 425–600 µm sterile glass beads (G9268, Sigma) and 3 ml of lysis buffer (10 mM Tris–HCl pH8, 25 mM Na2EDTA pH8, 1v/v% SDS and 100 mM NaCl). Samples were then subjected to bead beating, proteinase K (5 units/µL), and Lysozyme (2000 units/µL) digestion. All other DNA extraction steps were performed according to [23]. DNA was finally eluted in 40 µL EB (10 mM TE Tris 1 mM EDTA pH8). Wood samples were extracted using the DNeasy PowerWater Kit (Qiagen) according to the provided protocol to increase the DNA yield. DNA samples were kept at -20 °C for downstream applications.

### Fourier-transform infrared spectroscopy (FTIR)

Pieces of the plastic items (D1-D8) that were collected in the environment were also subjected to Fourier Transform Infrared (FTIR) analysis for the identification of their polymer composition (Supplementary Fig. S2). The FTIR was carried out using a JACSOFT/IR-6800 spectrometer. Spectra was obtained in the wavelength range of 200 cm ^-1^ to 4000 cm ^-1^, and the resolution was fixed at 2 cm ^-1^. A small plastic fragment of each sample was pressed under the apparatus K/Br crystal for measurements. The absorbance spectra were identified with the Open Specy open-source library [24].

### MinION library preparation and multiplexed nanopore sequencing

The sequencing runs were performed on three MinION flow cells in the main experiment and one flow cell in the second experiment, with 12-22 multiplexed libraries in each run. The libraries were prepared using the 16S barcoding kit (SQK-16S024, Oxford Nanopore Technologies) according to the manufacturer’s protocol. Ten ng of genomic DNA of each sample was prepared for PCR amplification with barcoded primers of the full-length 16S rRNA gene (27F: 5′-AGAGTTTGATCMTGGCTCAG3′ and 1492R: 5’-TACGGYTACCTTGTTACGACTT-3’). The barcoded libraries were pooled, loaded, and sequenced on MinION flow cells (FLO-MIN106D R9.4.1). Sequencing was carried out for 16-24 hours. Base-calling was done automatically by the MinKnow program. Raw reads were obtained in FAST5 and FASTQ formats from which “pass” quality reads were subjected to further analysis. All Nanopore MinION filtered reads analyzed in this project were deposited in the NCBI SRA database under BioProjects PRJNA1005105 and PRJNA1012293.

### Data analysis

The base-called reads were processed using Nanopore version 2.1.0, 16s workflow v2023.04.21-1804452 (https://epi2me.nanoporetech.com) on the EPI2ME desktop agent (ONT). The pipeline included sorting the reads into separate libraries according to the multiplexing barcodes, trimming the barcode and adaptor sequences, and comparing them to the NCBI bacterial 16S database. Reads < 800 bp or > 2000 bp were filtered out. The resulting CSV files from each run on EPI2ME were downloaded and further processed using R (V4.3.1). The processing steps included merging EPI2ME 16s CSV outputs of separate MinION runs into a single file (the main experiment), filtering out all reads that were not assigned to a barcode and reads that did not reach definitive classification (LCA≠0, either no classification or of more than one genus in the top three classifications). Lastly, multiplexing barcodes were converted into their corresponding samples, and read counts were normalized to relative abundances (percentages).

### Diversity analysis

Alpha diversity parameters were obtained using MicrobiomeAnalyst (Dhariwal, Chong et al. 2017) based on the taxonomic 16S metabarcoding datasets at the species level. Based on the mapped read counts of the samples (Supplementary Table S1) and the species richness rarefication curves (Supplementary Fig. S3A), we rarefied all samples to 7,940 mapped reads per sample. Community complexity within samples (alpha diversity) was estimated with Chao and Shannon indexes for richness and diversity accordingly. The analyses were performed according to the sample type (all time points combined) and according to the time points (all surfaces combined) in the main experiment. To assess the dissimilarity in the microbiome composition among the samples (beta diversity), we used Principal Component Analysis (PCA) and Non-Metric Multidimensional Scaling (NMDS) with Bray-Curtis Dissimilarity Distances. PCAtools package (R package version 2.12.0, https://github.com/kevinblighe/PCAtools) was used to produce the PCA plot. An *ANOVA* test with post hoc pairwise comparison tests was performed to identify significant differences between the groups.

### WGCNA analysis

The WGCNA analysis was performed using the WGCNA R package v1.71 (Langfelder and Horvath, 2008). Hierarchical clustering was then performed on the samples, and samples with a cut height below 7000 were retained (Supplementary Fig. S4A). Another filtering round was performed to remove bacteria with low variance (goodSamplesGenes (GSG) function within the WGCNA package). For the network topology analysis (TOM), soft-thresholding power β=10 was chosen (Supplementary Fig. S4B). First, a similarity matrix between each pair of bacteria across all samples was created. The similarity matrix was then transformed into an adjacency matrix, and the topological overlap matrix (TOM) along with the corresponding dissimilarity (1-TOM) value were calculated. Next, module eigengenes were calculated, and module-trait associations were quantified. Finally, the membership strength of individual species to their module and their significance to the relevant traits of that module were assessed. This allowed us to identify bacteria with relative preferences of surface type and time.

### Co-occurrence network visualization

The co-occurrence network analysis visualization was created using MicrobiomeAnalyst (Dhariwal, Chong et al. 2017) with low count filter parameters: minimum count – 20, mean abundance value – 30 (mbSet <-ApplyAbundanceFilter (mbSet, “mean”, 20, 0.3)) and low variance filter parameter: 10 % to remove (mbSet <-ApplyVarianceFilter (mbSet, “iqr”, 0.1)). Only plastic microbiome samples were used for the analysis using the SparCC algorithm [25] with a 0.05 p-value and 0.7 correlation threshold.

### Colonization assay with pure *Alcanivorax* strains

A total of five strains of the genus *Alcanivorax* were tested in this study, including: *Alcanivorax jadensis* (DSM 12178), *Alcanivorax dieselolei* (DSM 16502) and *Alcanivorax balearicus* (DSM 23776) from the DSMZ collection (Germany) and *Alcanivorax xenomutans* (KCTC 23751) and *Alcanivorax nanhaiticus* (KCTC 52137) from the Korean Collection for Type Cultures (KCTC, Korea). All stains were first cultivated in liquid or solid marine broth 2216 media (CAT: 76448, Sigma). The bacteria were cultured at 28° -30 °C until reaching the logarithmic phase (24-48 h, depending on the strain). Prior to the colonization experiment, bacteria cultures were pelleted by centrifugation at 5,000 g for 5 min and washed two times with carbon-free Bushnell-Haas basal mineral Media (BHM) (Bushnell and Haas, 1941) adjusted to pH 7.0 and supplemented with 30 g L^-1^ NaCl and 1 ml L^-1^ of trace metal solution (Wyman, Gregory et al. 1985) in double distilled water. To assess the effect of the different materials on the colonization rates, biofilm accumulation was quantified using crystal violet staining technique. The tested plastic surfaced included equal-size discs (made with a paper clipper), 0.5 mm in diameter and 0.15-0.6 mm-thickness of PE (IPETHENE® 323, Carmel olefins LTD), PP (CAPILENE® E 50 E, Carmel olefins LTD) and PET (SKYPET BR - Co-polymer PET, ResMart). Microscope glass coverslips (12 mm in diameter, Deckglaser) were used as the glass colonization surfaces. The plastic and glass disks underwent sterilization with 70% ethanol. The disks were placed in 50 ml flasks with 30 ml BHM. Each flask contained three technical replicates of each material. Each experiment was performed in three biological replicates (flasks). Plastic disks were fully submerged by securing them to each other with a fishing line. One ml of the tested *Alcanivorax* strains from the logarithmic growth phase (0.1 OD) was added to each of the experiment flasks. All flasks were incubated at 28° -30 °C with continuous agitation at 120 rpm for 48h. Control flasks contained the media without bacteria and were used as a blank for the crystal violet staining detection to ensure unspecific staining.

## Results

### Metabarcoding outcomes and microbiome diversity

A total of 58 samples were sequenced, of which 35 were from the main experiment, 15 from the second experiment, and eight from plastic debris (D1-D8). The average read number was 55,378 reads per sample (STDV = 32,465), of which 36% on average were mapped with high confidence to the database (with three top hits showing a single taxon, Supplementary Table S1). The average read size in all samples was 1491.5 nucleotides (SDV=23.5), the expected amplicon length representing the almost complete 16S rRNA gene sequence. The plastic microbiome samples had higher richness (*p*. value = 0.028) and biodiversity (*p*. value = 0.046) values compared to the glass samples but lower than the water samples (*p*. value = 1.931E-5 and 2.353E-5 accordingly) (Fig. 1A). No significant differences in the alpha diversity parameters were observed among the samples of the different newly-introduced plastic polymers (Supplementary Fig. S3B) nor between them and the debris samples. No significant differences were observed in the alpha diversity indexes between the different time points (all surfaces combined). However, the variance in the Shannon index values was smaller in the two days and 90 days time points compared to the 7, 14, and 30 days time points (Fig. 1A).

**Figure 1.**
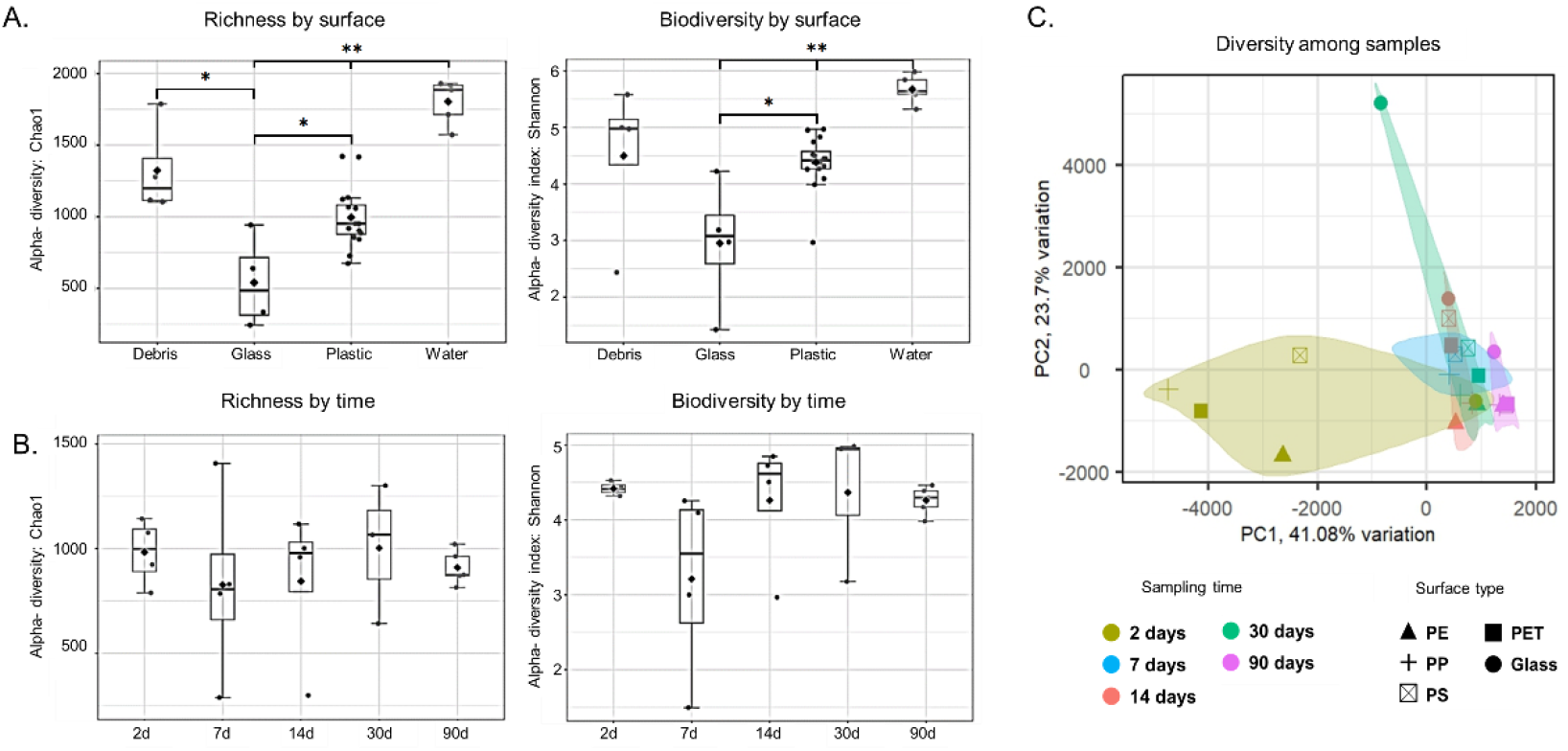
Diversity parameters. A. Alpha diversity indexes for richness (Chao1) and biodiversity (Shannon). B. PCA analysis for dissimilarity among samples (*ANOVA* test with post hoc pairwise comparison * *p* <0.05, ** *p* <0.01).

Dissimilarities in the microbiome taxonomic composition of the samples (beta diversity) were assessed using a PCA analysis (Fig. 1B). The samples in the PCA plot were partially clustered according to the time points but not according to the surface type. Based on principal component 1 in the PCA plot (PC1, X-axis, which explains 41.08% of the dissimilarities), the highest inter-sample diversity was between 2- and the 90-days samples, where the rest of the samples (7, 14 and 30 days) clustered together between them. The two days’ samples showed significantly higher dissimilarities among themselves than all other time points (Fig. 2B), suggesting a more substantial effect of the surface type in the early stages of biofilm development.

**Figure 2.**
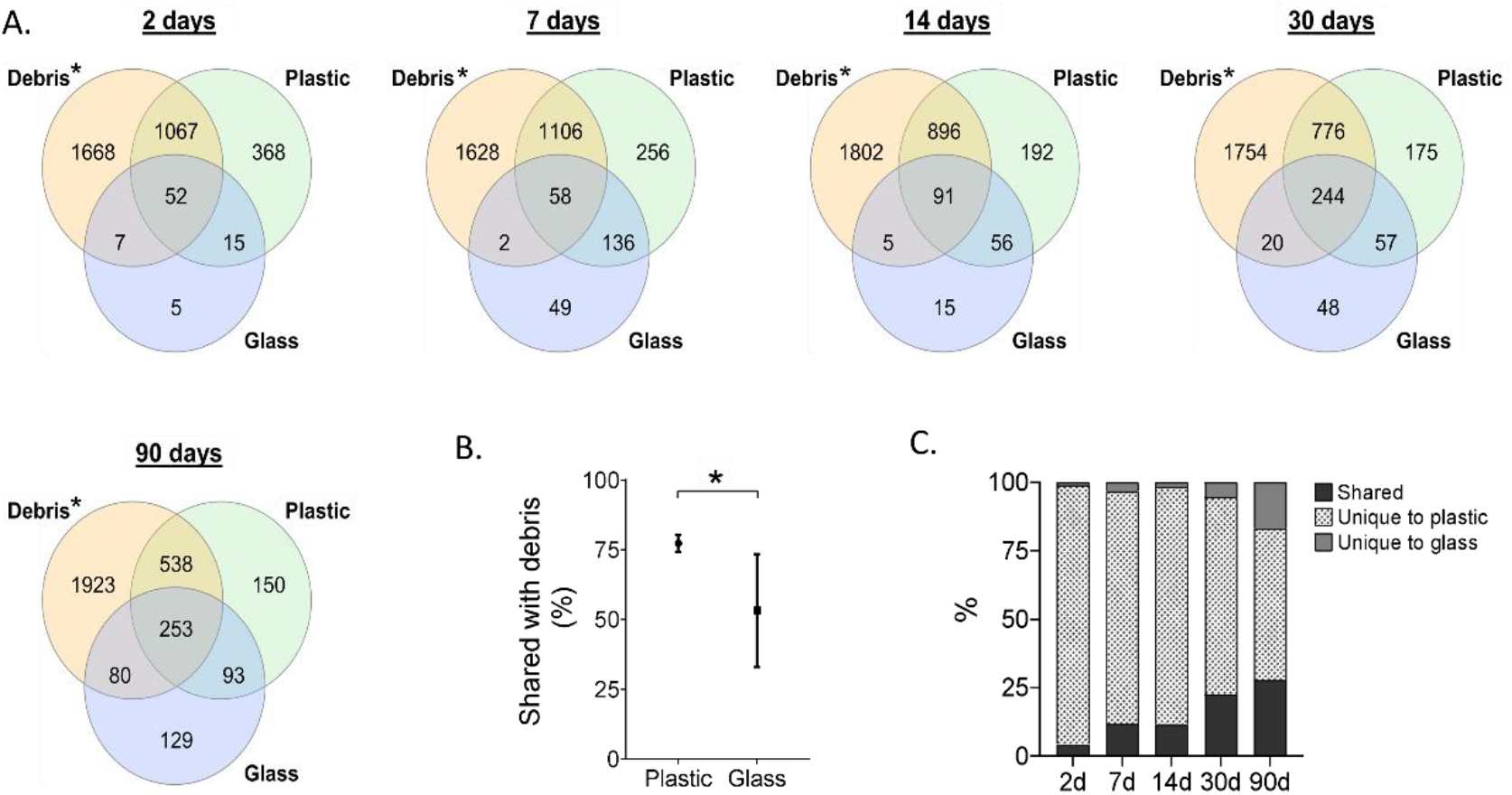
Shared and unique mapped species between debris, plastic (all polymers combined), and glass. A. Venn diagrams by time points. B. Percent of shared species between debris and plastic and between debris and glass (averages). C. Percent of shared and unique species between plastic and glass at each time point. * Refer to the initial debris microbiome composition (at the beginning of the experiment).

### Plastic-attached microbiome becomes less specific with time

A versatile mixture of different plastic items was introduced in the experiment to obtain a diverse repertoire of plastic-associated bacteria for the inoculation of the new surfaces. The debris included fragments or complete items, including cups, bags, straws, plastic cutlery, bottles, and several other, mostly fragmented, plastic items. An FTIR analysis that was performed on pieces of the four items of each of the two aquarium experiments (D1-D8) showed that the items contained different plastic polymers, including (but not limited to) PE, PP, and PET (Supplementary Fig. S2).

To assess the proportions of bacteria that migrated to the newly introduced surfaces, we compared the total number of the mapped species of the sampled debris (D1-D4 for the main experiment and D5-D8 in the second experiment) with that of all plastic surfaces in each time point. The results of the main experiment suggested that 74.5% of the mapped species were shared between the debris and the plastic surfaces as soon as two days after the beginning of the experiment, indicating fast bacterial migration from the debris to the surfaces. This proportion stayed within the 68.4-78.1% range throughout the experiment until 90 days (Fig. 2A). These numbers may be underestimated, given that not all debris was sampled. On the other hand, as expected, the proportions of the shared mapped species between the glass surfaces and the debris were significantly lower (Fig. 2B). When comparing the mapped species lists of plastics vs. glass, we identified a persistent trend of increase in the proportions of the shared species between the two surface types (Fig. 2C), indicating a gradual loss of microbiome surface-specificity.

### The abundance dynamics of dominant genera on different plastic polymers are similar under different temperatures

Identifying dominant taxa within a specific sample is often done based on its high read proportions within the total read count for this sample. Here, the top six most abundant genera within the pooled microbiome of each polymer (i.e. from all the time points for that polymer) were analyzed (Fig. 3). Most (9/16) of the identified dominant genera were shared between the main and the second experiment. The shared genera included *Alcanivorax, Alteromonas, Ketobacter, Pseudomonas, Porticoccus, Marinobacter, Sneathiella, Microbulbifer* and *Spongiobacter*. Moreover, the abundance dynamics of these genera appeared to be similar in most cases between the two experiments - *Alcanivorax, Ketobacter, Pseudomonas, Porticoccus, Sneathiella, Microbulbifer*, and *Spongiobacter* were all more abundant in PE compared to PET, glass and wood between 7- and 30-days’ time points and *Alteromonas* seems to be an early generalist colonizer in both experiments.

**Figure 3.**
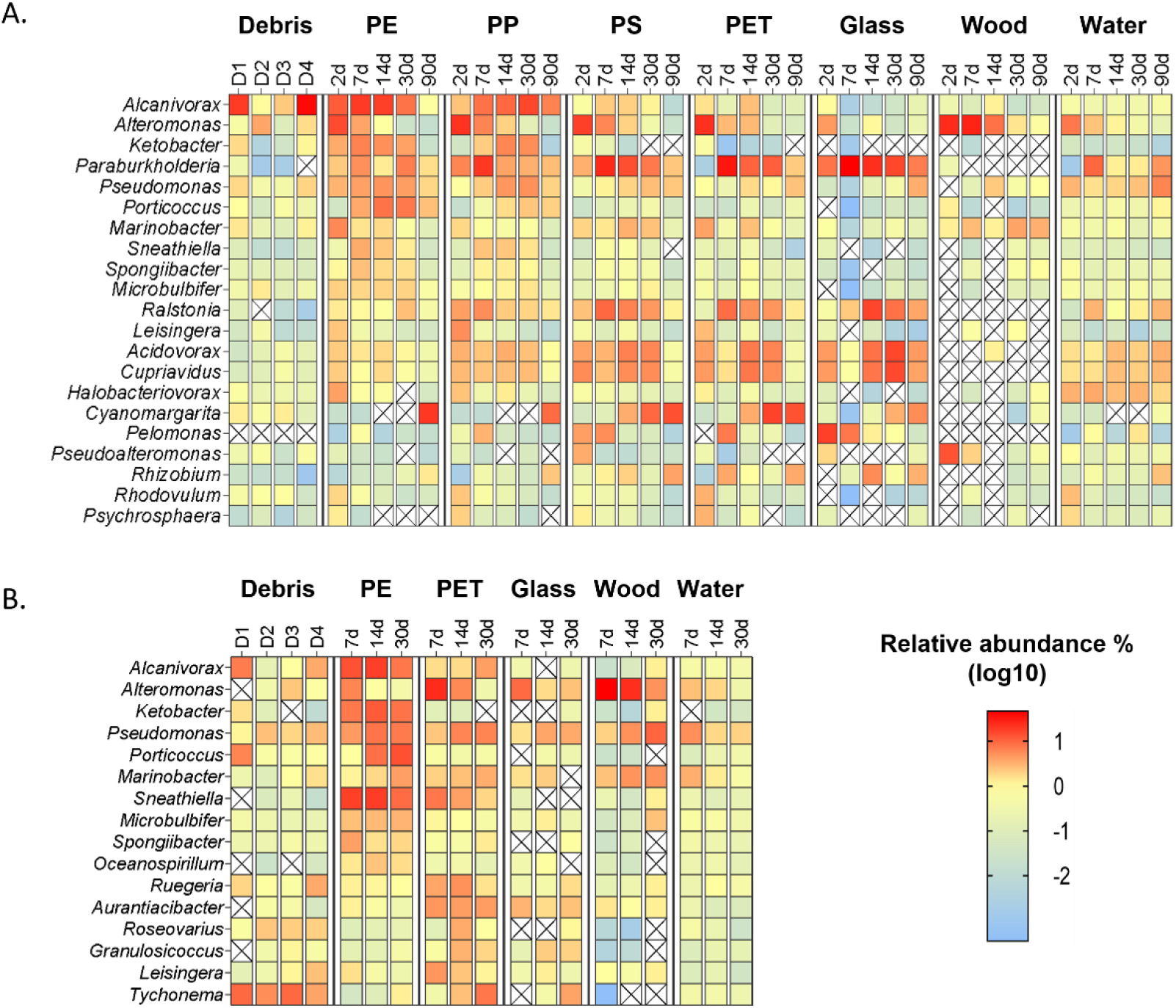
Relative abundance of top abundant genera. A. first (primary) experiment at 28±2 °C. B. second experiment at 35±3 °C. The top 6 genera of each of the plastic polymer types are represented.

### Identification of key genera within succession modules and genera with general polymer-specific preference using WGCNA

Weighted gene co-expression network analysis (WGCNA) is used to identify correlations among genes based on their expression patterns. Here, we implemented WGCNA to identify correlations among bacteria based on their projected relative abundances. A signed co-occurrence network was created for the main experiment. Clusters of the mapped bacterial species were detected by hierarchical clustering based on their read counts in each surface and time point (Fig. 4A). The clustering was used to create module eigengenes. To reduce the number of modules, a merged dynamic analysis was performed (Fig. 4B), resulting in a total of 26 modules (marked in colors, Fig 4C) with module sizes ranging from 36 species (dark orange) to 541 species (Turquoise). A module-trait relationship table was created based on the module eigengenes. Most module eigengenes had a positive (often significant) correlation to a single polymer and a specific time point. For example, the Green Yellow module (Fig 4C, first raw) positively correlated to glass and 30d traits (c = 0.5, *p* = 0.02 and c = 0.37, *p* = 0.09, respectively), while it was negatively correlated to all other traits.

**Figure 4.**
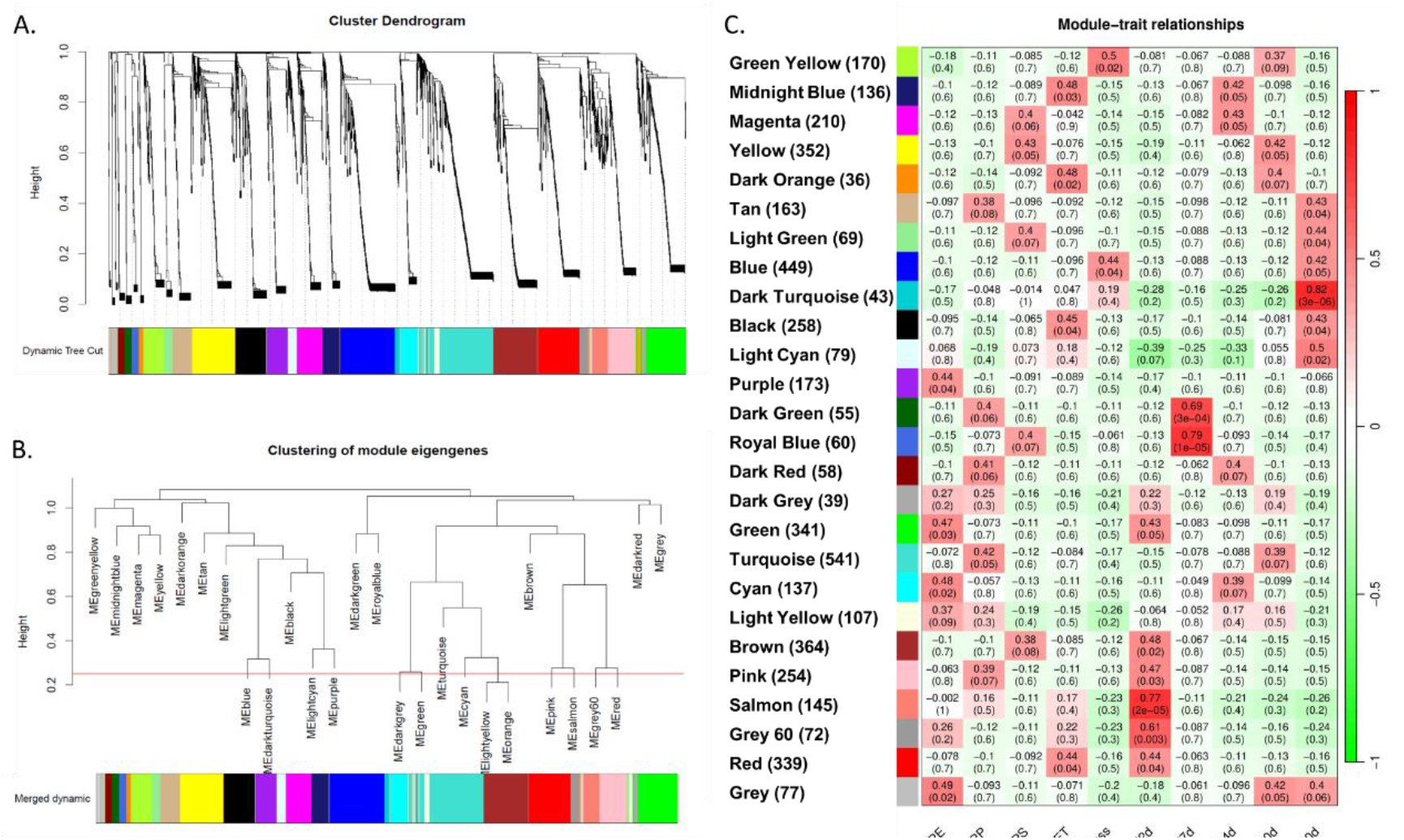
Cluster dendrograms and module assignments. A. cluster dendrogram based on the relative abundance patterns of identified bacterial species. Hierarchical clustering was done using topology overlap measure (TOM). The colours represent the module assignments of the clusters. B. merged dynamic module clustering of similar eigengenes based on a merging threshold 0.25. C. module eigengenes table with module-trait corresponding correlation and significance values.

To identify key genera within specific modules, we looked for species with a high correlation to the module eigengenes (module membership) and trait (surface type/time). A key genus in a module was defined as a genus containing at least three species with > 0.9 module membership score. For example, *Alteromonas* was identified as a key genus in the Salmon module, as it contained 11 mapped species with > 0.9 correlation to the module (Supplementary Fig. S5A). Similar to the module eigengene, this genus was not significant to a specific surface but correlated significantly to the 2d time point. Nevertheless, most of the *Alteromonas* species in that module showed a negative correlation to the glass and a positive correlation to the plastic surfaces (Supplementary Fig. S5B). These results imply that these *Alteromonas* bacteria are early colonizers of different plastic polymers. Another example is the genus *Dokdonia*, which had six mapped species in the brown module (Supplementary Fig. S5C) with a strong positive correlation to the PS surface and 2d-time point (Supplementary Fig. S5D), suggesting they are early colonizers on PS. Genus *Nevskia*, on the other hand (with six mapped species in the Turquoise module; Supplementary Fig. S5E), had a strong positive correlation to PP and 30d (Supplementary Fig. S5F), suggesting it is a late colonizer on this polymer. Some genera were identified as key genera in multiple modules. For example, the genus *Pseudomonas* was identified as a key genus in five different modules, especially in the green module (30 in total, of which 10 with module membership > 0.9) and the turquoise module (43 mapped species in total, of which 9 with module membership > 0.9). Accordingly, the genus *Pseudomonas* showed a high correlation to PE and 2d as well as to PP and 30d (Supplementary Fig. S6). These results suggest that the genus *Pseudomonas* contain several subgroups with different succession patterns. Using this filtration threshold, we identified 77 module key genera within 21 modules (summarized in Table 1), of which 17 modules positively correlated to a single specific surface type and time-point. Interestingly, glass-related key genera within the green-yellow and blue modules were correlated with the later succession stages (at the 30d and mainly 90d time points) and were missing from the early time points (Table 1).

**Table 1:**
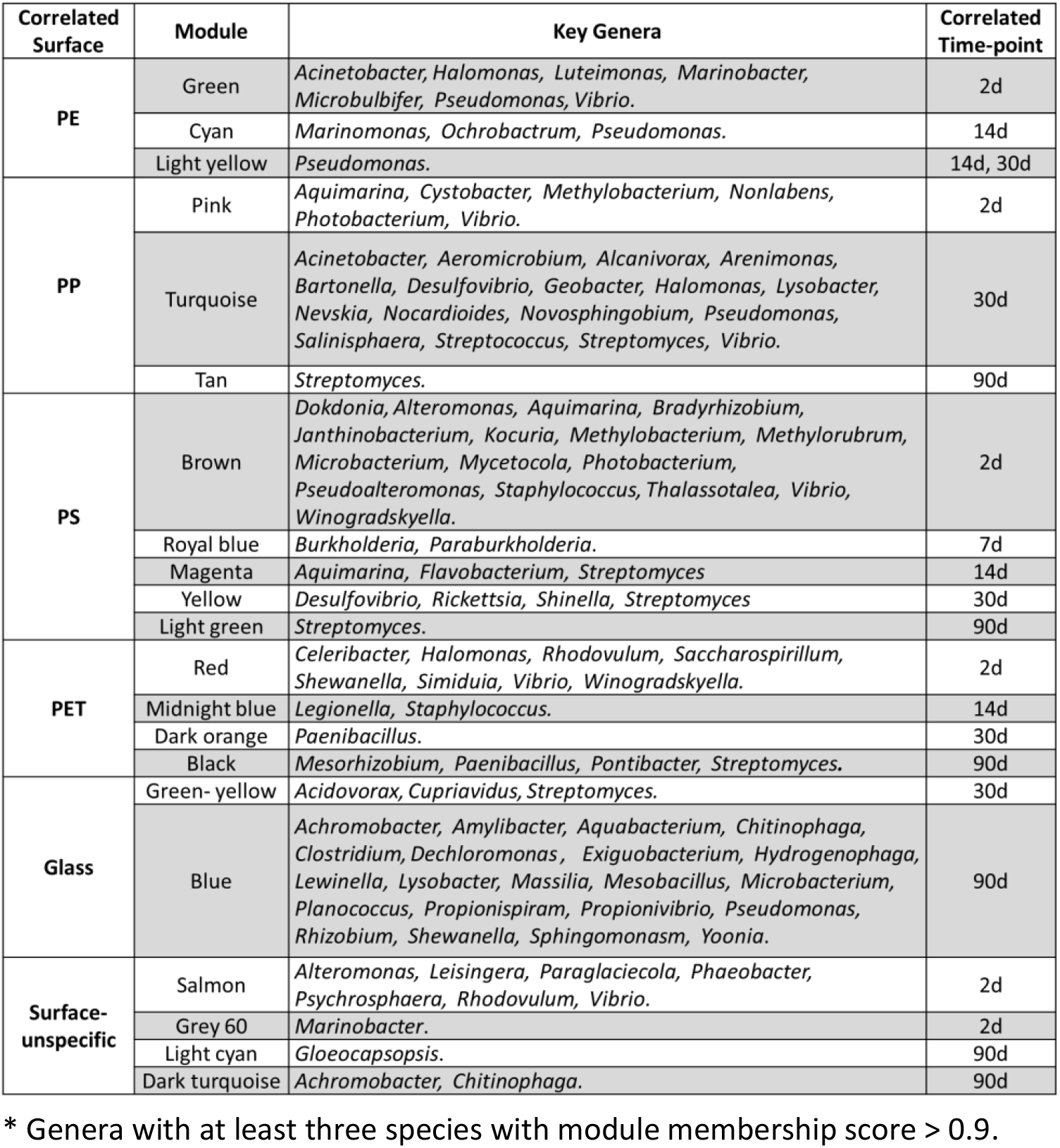
Key genera by modules*.

To identify genera enriched on specific plastic polymers, we screened for species with significant correlations to surface type traits compared to the other surfaces within the first month of the experiment (including 2d, 7d, 14d, and 30d time points). We selected genera with gene significant to surface *p*. value ≤ 0.05 and a relative abundance of more than 0.1%. Using this filtration threshold, we identified 39 genera with potential significant enrichment to specific plastic polymers (13 genera for PE, 11 for PP, 9 for PS, and 6 for PET; Table 2).

**Table 2.**
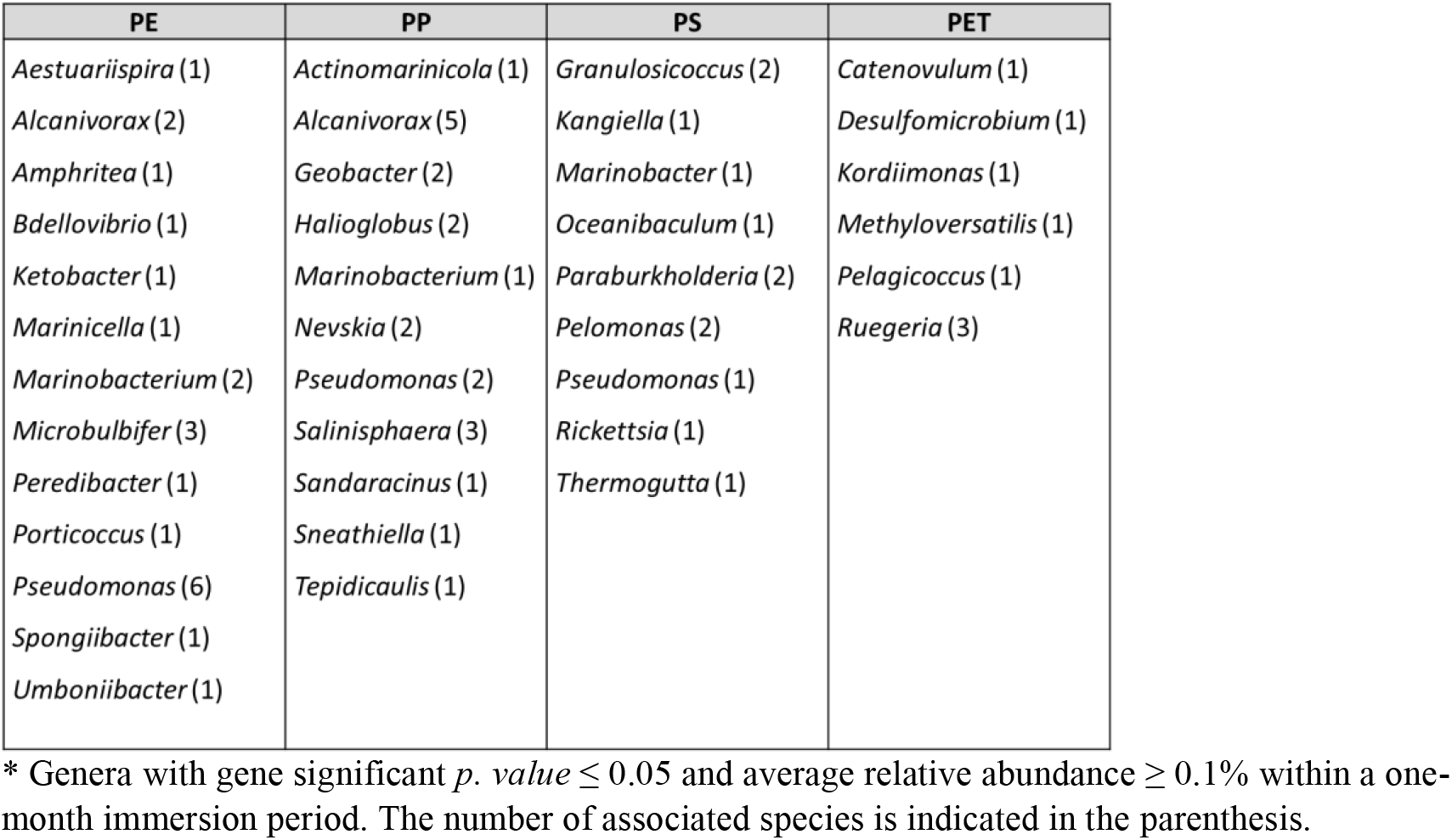
Significantly enriched genera on plastic polymers *.

### The co-occurrence network shows dominant genera segregation according to their polymer preferences

To better understand the relationships among different bacteria on the plastic surfaces, we investigated the intra-genus connections within the co-occurrence network while applying minimum threshold values for genus abundance, diversity among samples and connection strength (Fig. 5).

**Figure 5.**
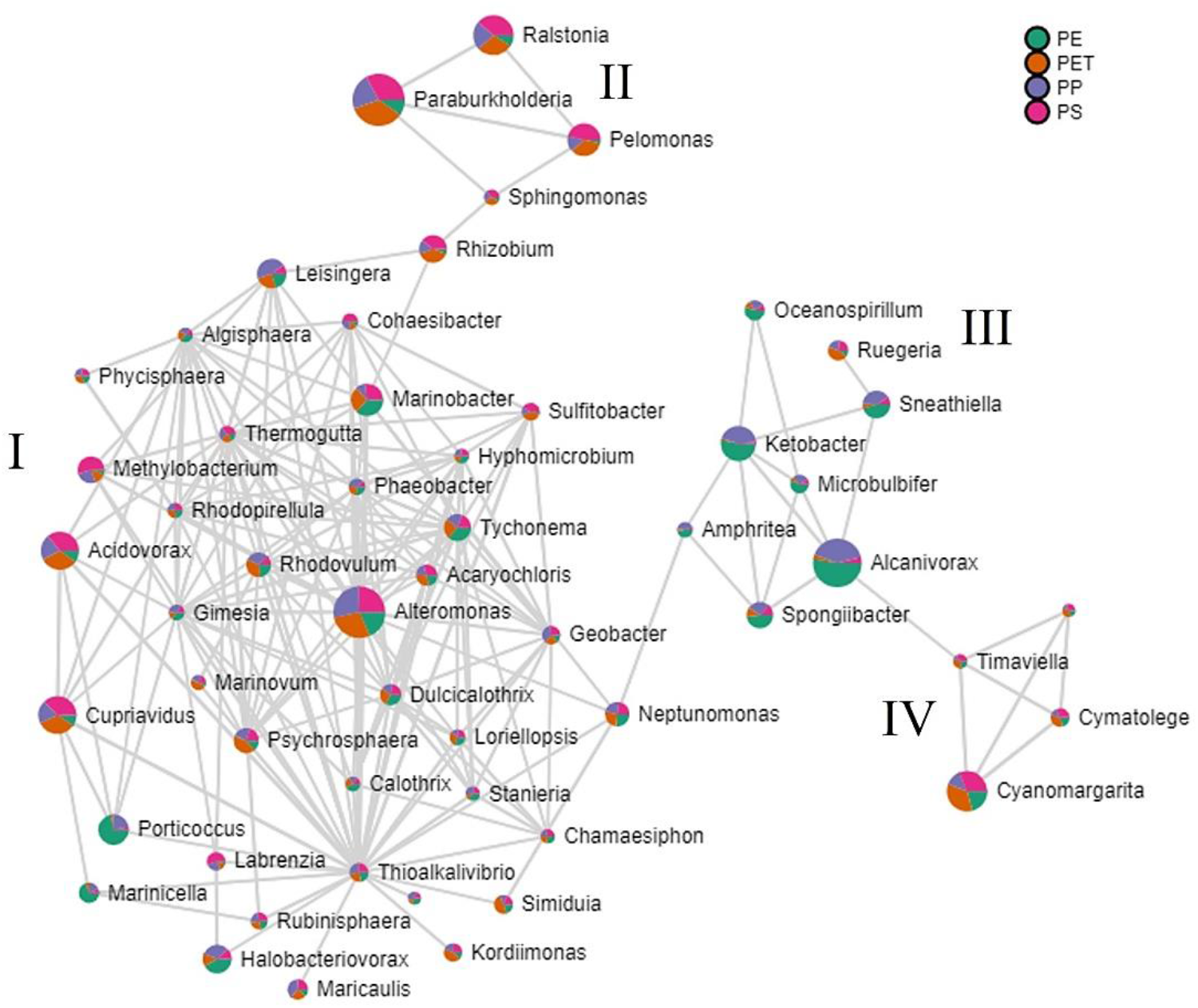
co-occurrence network of dominant plastisphere genera. Only genera that passed the abundance filter are represented (see methods section for details). The relative abundances of the represented genera on the different plastic polymers are shown in the pie charts in different colours: Green - PE, purple - PP, PET - orange and PS - pink. The pie chart sizes represent the general prevalence of each genus. Four identified clusters are marked with Roman letters (I-IV)

The network visualization resulted in four partially segregated clusters. The major cluster (cluster “I”) contained most of the genera, and the three smaller ones (clusters II, III, IV) included the rest. While cluster I contained genera of mixed polymer-based abundance signatures, clusters II, III and IV appeared to contain genera with almost unified signatures. Cluster III contained seven genera significantly enriched on polyolefins (PE and PP), with *Alcanivorax* at the top, followed by *Ketobacter* and *Sneathiella*. Cluster II contained four genera with a relatively higher abundance of PS and PET followed by PP, with *Paraburkholderia* being the most abundant in this cluster. Cluster IV contained four cyanobacteria genera with higher abundance on PS and PET, followed by PE. The dominant genus in this cluster was *Cyanomargarita*.

### Testing the succession dynamics of bacteria from the genus *Alcanivorax* in pure cultures

*The alcanivorax* genus was the most dominant on PE and PP (Fig. 3, Fig. 5) and was found to be significantly enriched on both polymer types (Table 2). The reads mapped to *Alcanivorax* species accounted for 29% of the total PE microbiome mapped reads count after 14 days and 23% of the total PP microbiome mapped reads count after 30 days. Seven bacteria strains that have been mapped by our metabarcoding pipeline to *Alcanivorax* species showed higher relative abundance on polyolefin (PE and PP) surfaces over other surfaces (Fig. 6A). WGCNA for these strains resulted in two main subgroups according to their succession patterns. Strains that were mapped to species *A. Jedensis, A. nanhaticus* and *A. balericus* showed positive abundance correlation to PE and 2d, while those that have been mapped to *A. dieselolei, A. gelatiniphagus, A. hongdegensis, A. marinus*, and *A. xenomutans* showed positive abundance correlation to PP and 30d (Fig. 6B). We followed up these results performing colonization experiments with pure cultures of five of the identified *Alcanivorax* species that were purchased for this purpose. The colonization of each of the purchased *Alcanivorax* species on PE, PP, PET, and glass was tested after two days of immersion using crystal violet staining method (Fig. 6C, left). The results showed a significant preference for colonizing polyolefins over PET and glass in *A. balearicus, A. diselolei*, and *A. xenomutans*, while *A. nanhaticus* and *A. Jedensis* did not show adhesion to any of the surfaces (Fig. 6C, right).

**Figure 6.**
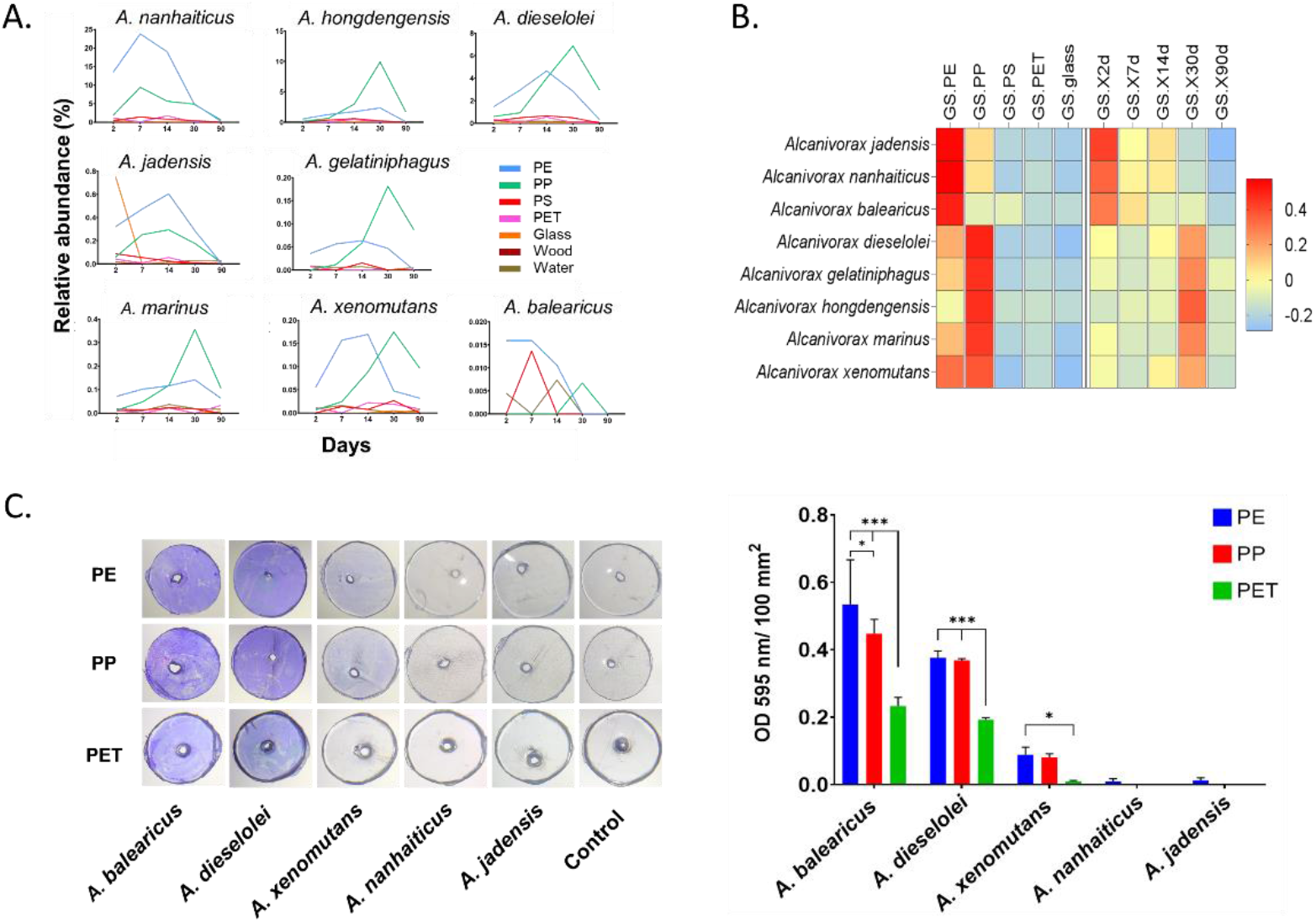
Detailed relative abundance dynamics and plastic colonization preference of *Alcanivorax* bacteria. A. Relative abundance of *Alcanivorax* species on surfaces during experiment time points B. The corresponding gene significance (GS) charts depict two sub-groups according to the level of correlation to the different traits (surface types and time points). C. biofilm formation of *Alcanivorax* bacteria by crystal violet staining after 48h of immersion. Left - Photos of the discs after staining and washing steps, Right - Biofilm quantification 595 nm (Two-way ANOVA test, * *P* <0.05, *** *P* <0.01).

## Discussion

Plastic debris in the ocean is made of versatile combinations of different polymers and additives, providing them with different chemical and physical properties. Very little is known about how these properties affect the colonization and succession of microbial species. Here, we tested the successive development of bacterial microbiome on four of the most abundant plastic polymer pollutants in marine environments, namely, PE, PP, PS, and PET. These polymers are different in their chemical structure; PE, PP, and PS consist of a C-C backbone with different side groups (hydrogen, methyl, and phenyl ring, respectively), while PET is a co-polymer of ethylene glycol and terephthalic acid with an ester group in its backbone. Those chemical differences give each polymer different properties, including density, hydrophobicity, roughness, surface energy, and chemical reactivity that affect the adherence and growth of marine microorganisms.

It was shown that as the plastic biofilm cover develops, the effect of the polymer type on the microbiome composition is weakening until it is almost unnoticeable [2, 17]. This notion was validated in a recent study that showed that the polymer type affects microbiome composition and metabolic functionality in early marine plastic communities but not mature communities [26]. Using linear regression analysis, we recently identified prokaryotic and eukaryotic taxa that were significantly enriched on specific plastic polymers in the marine environment [20]. However, in agreement with the previous studies mentioned above, we showed here that greater variability is expected during even earlier developmental stages of the plastic communities, as the different polymer samples were most dissimilar in the 2-day time point, followed by a gradual, consistent shift toward equilibrium. This notion was also supported by the constant increase in the portion of the shared species between the plastic and the glass surfaces with time.

To better understand the plastisphere bacterial microbiome development on different plastic and non-plastic surfaces, we first screened our long-read 16S rRNA metabarcoding databases for dominant genera based on their relative abundance on specific surface type and time. This analysis provides important data on key genera colonizing the surfaces but without statistical significance validation. When comparing the lists of the key genera of the main experiment versus those of the second experiment, we found that in spite of the different temperature ranges (7°C on average), the different content of the debris brought from the environment, and the smaller polymer variety and time range in the second experiment, most of the top-abundant genera (9/16) in the second experiment were shared with those of the main experiment, suggesting that the polymer-based preference mechanisms are highly robust and less effected from environmental factors.

Next, we used WGCNA as a tool to analyze the co-occurrence network of bacterial species within the microbiome of the different surfaces. WGCNA was coopted before for similar purposes [27-29]. The implementation of WGCNA in this study allowed us to obtain, for the first time, succession signatures of the plastisphere bacteria at the genus and species levels and to understand their relationship to the surface type and time-point. The validity of our WGCNA results was best supported by the fact that multiple bacteria that were mapped to a specific genus were clustered in the specific module/s, with a high correlation to their module eigengene in most cases. The results of this study are also supported by earlier studies. For example, in agreement with our results, genus *Alteromonas* was found to consist of pioneer generalist colonizer species (Dang and Lovell 2000, Nam et al. 2008, Pinto, Langer et al. 2019).

Moreover, several bacterial genera, including *Marinobacter* and *Marinobacterium*, that were significantly enriched on plastic polymers in this study are known hydrocarbon metabolizers [30, 31]. Furthermore, at least three of the genera that were significantly enriched on PE, according to our analysis, were previously shown to degrade PE, including *Alcanivorax* [32], *Microbulbifer* [33] and *Pseudomonas* [34]. Using the microbiomeanalyst platform, we identified four co-occurrence clusters of dominant bacteria, of which 3 clusters contained genera with similar polymer-based occurrence signatures. The tight connections among genera within each cluster may imply tight collaborative interaction.

The simple experiment design presented in this study, together with the implementation of the MinION nanopore-based metabarcoding pipeline and the analysis tools that were coopted here, may be practical and applicable in other fields of microbiome research. For example, it may be used in the biomedical field to understand the colonization and succession of pathogenic bacteria on artificial implants. Further calibration of the different parameters in the analyses will be needed to address such future research questions. Nevertheless, we see great potential in the research approach presented here to understand better microbial succession processes as well as preference for specific surfaces or times.

## References

1. Amaral-Zettler, L. A., et al. (2021). “Diversity and predicted inter-and intra-domain interactions in the Mediterranean Plastisphere.” Environmental Pollution 286: 117439.

2. Amaral-Zettler, L. A., et al. (2020). “Ecology of the Plastisphere.” Nature Reviews Microbiology 18(3): 139–151.

3. Bos, R. P., et al. (2023). “Plastics select for distinct early colonizing microbial populations with reproducible traits across environmental gradients.” Environmental microbiology.

4. Bushnell, L. and H. Haas (1941). “The utilization of certain hydrocarbons by microorganisms.” Journal of bacteriology 41(5): 653–673.

5. Coons, A. K., et al. (2021). “Biogeography rather than substrate type determines bacterial colonization dynamics of marine plastics.” PeerJ 9: e12135.

6. Cowger, W., et al. (2021). “Microplastic spectral classification needs an open source community: open specy to the rescue!” Analytical Chemistry 93(21): 7543–7548.

7. Debeljak, P., et al. (2017). “Extracting DNA from ocean microplastics: a method comparison study.” Analytical methods 9(9): 1521–1526.

8. Dhariwal, A., et al. (2017). “MicrobiomeAnalyst: a web-based tool for comprehensive statistical, visual and meta-analysis of microbiome data.” Nucleic acids research 45(W1): W180–W188.

9. Duan, J., et al. (2021). “Weathering of microplastics and interaction with other coexisting constituents in terrestrial and aquatic environments.” Water Research 196: 117011.

10. Langfelder, P. and S. Horvath (2008). “WGCNA: an R package for weighted correlation network analysis.” BMC bioinformatics 9(1): 1–13.

11. Latva, M., et al. (2022). “Microbial pioneers of plastic colonization in coastal seawaters.” Marine Pollution Bulletin 179: 113701.

12. Li, J., et al. (2020). “Are bacterial communities associated with microplastics influenced by marine habitats?” Science of The Total Environment 733: 139400.

13. Marsay, K. S., et al. (2022). “High-resolution screening for marine prokaryotes and eukaryotes with selective preference for polyethylene and polyethylene terephthalate surfaces.” Frontiers in Microbiology 13: 845144.

14. Oberbeckmann, S., et al. (2014). “Spatial and seasonal variation in diversity and structure of microbial biofilms on marine plastics in Northern European waters.” FEMS microbiology ecology 90(2): 478–492.

15. Oberbeckmann, S., et al. (2016). “Microbes on a bottle: substrate, season and geography influence community composition of microbes colonizing marine plastic debris.” PloS one 11(8): e0159289.

16. Ogonowski, M., et al. (2018). “Evidence for selective bacterial community structuring on microplastics.” Environmental microbiology 20(8): 2796–2808.

17. Sérvulo, T., et al. (2023). “Plastisphere composition in a subtropical estuary: Influence of season, incubation time and polymer type on plastic biofouling.” Environmental Pollution 332: 121873.

18. Wallbank, J. A., et al. (2022). “Into the Plastisphere, where only the generalists thrive: early insights in plastisphere microbial community succession.” Frontiers in Marine Science 9: 841142.

19. Ward, C. S., et al. (2022). “Microbiome development of seawater-incubated pre-production plastic pellets reveals distinct and predictive community compositions.” Frontiers in Marine Science 8: 807327.

20. Wyman, M., et al. (1985). “Novel role for phycoerythrin in a marine cyanobacterium, Synechococcus strain DC2.” Science 230(4727): 818–820.

21. Xu, X., et al. (2019). “Marine microplastic-associated bacterial community succession in response to geography, exposure time, and plastic type in China’s coastal seawaters.” Marine Pollution Bulletin 145: 278–286.

22. Zettler, E. R., et al. (2013). “Life in the “plastisphere”: microbial communities on plastic marine debris.” Environmental science & technology 47(13): 7137–7146.

23. Amaral-Zettler, L. A., et al. (2021). “Diversity and predicted inter-and intra-domain interactions in the Mediterranean Plastisphere.” Environmental Pollution 286: 117439.

24. Amaral-Zettler, L. A., et al. (2020). “Ecology of the plastisphere.” Nature Reviews Microbiology 18(3): 139–151.

25. Dang, H. and C. R. Lovell (2000). “Bacterial primary colonization and early succession on surfaces in marine waters as determined by amplified rRNA gene restriction analysis and sequence analysis of 16S rRNA genes.” Applied and environmental microbiology 66(2): 467–475.

26. Duan, J., et al. (2021). “Weathering of microplastics and interaction with other coexisting constituents in terrestrial and aquatic environments.” Water Research 196: 117011.

27. Erni-Cassola, G., et al. (2020). “Early colonization of weathered polyethylene by distinct bacteria in marine coastal seawater.” Microbial ecology 79: 517–526.

28. Hansen, J., et al. (2021). “Effect of polymer type on the colonization of plastic pellets by marine bacteria.” FEMS microbiology letters 368(5): fnab026.

29. Lee, J.-W., et al. (2008). “Bacterial communities in the initial stage of marine biofilm formation on artificial surfaces.” The journal of microbiology 46: 174–182.

30. Marsay, K. S., et al. (2023). “The geographical and seasonal effects on the composition of marine microplastic and its microbial communities: The case study of Israel and Portugal.” Frontiers in Microbiology 14: 1089926.

31. Oberbeckmann, S., et al. (2018). “Environmental factors support the formation of specific bacterial assemblages on microplastics.” Frontiers in Microbiology 8: 2709.

32. Oberbeckmann, S., et al. (2014). “Spatial and seasonal variation in diversity and structure of microbial biofilms on marine plastics in Northern European waters.” FEMS microbiology ecology 90(2): 478–492.

33. Oberbeckmann, S., et al. (2016). “Microbes on a bottle: substrate, season and geography influence community composition of microbes colonizing marine plastic debris.” PloS one 11(8): e0159289.

34. Pinto, M., et al. (2019). “The composition of bacterial communities associated with plastic biofilms differs between different polymers and stages of biofilm succession.” PloS one 14(6): e0217165.

35. Ward, C. S., et al. (2022). “Microbiome development of seawater-incubated pre-production plastic pellets reveals distinct and predictive community compositions.” Frontiers in Marine Science 8: 807327.

36. Zettler, E. R., et al. (2013). “Life in the “plastisphere”: microbial communities on plastic marine debris.” Environmental science & technology 47(13): 7137–7146.

1. Latva, M., et al., Microbial pioneers of plastic colonization in coastal seawaters. Marine Pollution Bulletin, 2022. 179: p. 113701.

2. Amaral-Zettler, L.A., E.R. Zettler, and T.J. Mincer, Ecology of the plastisphere. Nature Reviews Microbiology, 2020. 18(3): p. 139–151.

3. Zettler, E.R., T.J. Mincer, and L.A. Amaral-Zettler, Life in the “plastisphere”: microbial communities on plastic marine debris. Environmental Science & Technology, 2013. 47(13): p. 7137–7146.

4. Oberbeckmann, S., et al., Spatial and seasonal variation in diversity and structure of microbial biofilms on marine plastics in Northern European waters. FEMS microbiology ecology, 2014. 90(2): p. 478–492.

5. Oberbeckmann, S., A.M. Osborn, and M.B. Duhaime, Microbes on a bottle: substrate, season and geography influence community composition of microbes colonizing marine plastic debris. PLoS One, 2016. 11(8): p. e0159289.

6. Amaral-Zettler, L.A., et al., diversity and predicted inter-and intra-domain interactions in the Mediterranean Plastisphere. Environmental Pollution, 2021. 286: p. 117439.

7. Marsay, K.S., et al., The geographical and seasonal effects on the composition of marine microplastic and its microbial communities: The case study of Israel and Portugal. Frontiers in Microbiology, 2023. 14: p. 1089926.

8. Xu, X., et al., Marine microplastic-associated bacterial community succession in response to geography, exposure time, and plastic type in China’s coastal seawaters. Marine pollution bulletin, 2019. 145: p. 278–286.

9. Wallbank, J.A., et al., Into the Plastisphere, Where Only the Generalists Thrive: Early Insights in Plastisphere Microbial Community Succession. Frontiers in Marine Science, 2022: p. 626.

10. Oberbeckmann, S., B. Kreikemeyer, and M. Labrenz, Environmental factors support the formation of specific bacterial assemblages on microplastics. Frontiers in microbiology, 2018. 8: p. 2709.

11. Erni-Cassola, G., et al., Early colonization of weathered polyethylene by distinct bacteria in marine coastal seawater. Microbial ecology, 2020. 79(3): p. 517–526.

12. Duan, J., et al., Weathering of microplastics and interaction with other coexisting constituents in terrestrial and aquatic environments. Water Research, 2021. 196: p. 117011.

13. Dussud, C., et al., Evidence of niche partitioning among bacteria living on plastics, organic particles and surrounding seawaters. Environmental Pollution, 2018. 236: p. 807–816.

14. Li, J., et al., Are bacterial communities associated with microplastics influenced by marine habitats? Science of The Total Environment, 2020. 733: p. 139400.

15. Coons, A.K., et al., Biogeography rather than substrate type determines bacterial colonization dynamics of marine plastics. PeerJ, 2021. 9: p. e12135.

16. Sérvulo, T., et al., Plastisphere composition in a subtropical estuary: Influence of season, incubation time and polymer type on plastic biofouling. Environmental Pollution, 2023. 332: p. 121873.

17. Pinto, M., et al., The composition of bacterial communities associated with plastic biofilms differs between different polymers and stages of biofilm succession. PloS one, 2019. 14(6): p. e0217165.

18. Hansen, J., et al., Effect of polymer type on the colonization of plastic pellets by marine bacteria. FEMS microbiology letters, 2021. 368(5): p. fnab026.

19. Ward, C.S., et al., Microbiome Development of Seawater-Incubated Pre-Production Plastic Pellets Reveals Distinct and Predictive Community Compositions. Frontiers in Marine Science, 2022.

20. Marsay, K., et al., High-resolution screening for marine prokaryotic and eukaryotic taxa with selective preference for PE and PET surfaces. Frontiers in microbiology 2021.

21. Yokoyama, D., et al., Large-scale omics dataset of polymer degradation provides robust interpretation for microbial niche and succession on different plastisphere. ISME communications, 2023. 3(1): p. 67.

22. Silva, V., V. Pérez, and B.M. Gillanders, Short-term plastisphere colonization dynamics across six plastic types. Environmental Microbiology, 2023.

23. Debeljak, P., et al., Extracting DNA from ocean microplastics: a method comparison study. Analytical methods, 2017. 9(9): p. 1521–1526.

24. Cowger, W., et al., Microplastic spectral classification needs an open source community: open specy to the rescue! Analytical Chemistry, 2021. 93(21): p. 7543–7548.

25. Friedman, J. and E.J. Alm, Inferring correlation networks from genomic survey data. PLoS computational biology, 2012. 8(9): p. e1002687.

26. Bos, R.P., et al., Plastics select for distinct early colonizing microbial populations with reproducible traits across environmental gradients. Environmental Microbiology, 2023.

27. Djurhuus, A., et al., Environmental DNA reveals seasonal shifts and potential interactions in a marine community. Nature communications, 2020. 11(1): p. 254.

28. Chen, H., et al., Bacterial community response to chronic heavy metal contamination in marine sediments of the East China Sea. Environmental Pollution, 2022. 307: p. 119280.

29. Li, Q., et al., Soil Geochemical Properties Influencing the Diversity of Bacteria and Archaea in Soils of the Kitezh Lake Area, Antarctica. Biology, 2022. 11(12): p. 1855.

30. Gauthier, M.J., et al., Marinobacter hydrocarbonoclasticus gen. nov., sp. nov., a new, extremely halotolerant, hydrocarbon-degrading marine bacterium. International Journal of Systematic and Evolutionary Microbiology, 1992. 42(4): p. 568–576.

31. Bae, S.S., et al., Marinobacterium aestuarii sp. nov., a benzene-degrading marine bacterium isolated from estuary sediment. International Journal of Systematic and Evolutionary Microbiology, 2018. 68(2): p. 651–656.

32. Zadjelovic, V., et al., A Mechanistic Understanding of Polyethylene Biodegradation by the Marine Bacterium Alcanivorax. Journal of Hazardous Materials, 2022: p. 129278.

33. Li, Z., et al., Biodegradation of low-density polyethylene by Microbulbifer hydrolyticus IRE-31. Journal of environmental management, 2020. 263: p. 110402.

34. Kyaw, B.M., et al., Biodegradation of low density polythene (LDPE) by Pseudomonas species. Indian journal of microbiology, 2012. 52: p. 411–419.

