## Supplemental for "Differential colonization and succession dynamics of marine bacteria on different plastic polymers"

* Equal authorship

### Supplementary Results

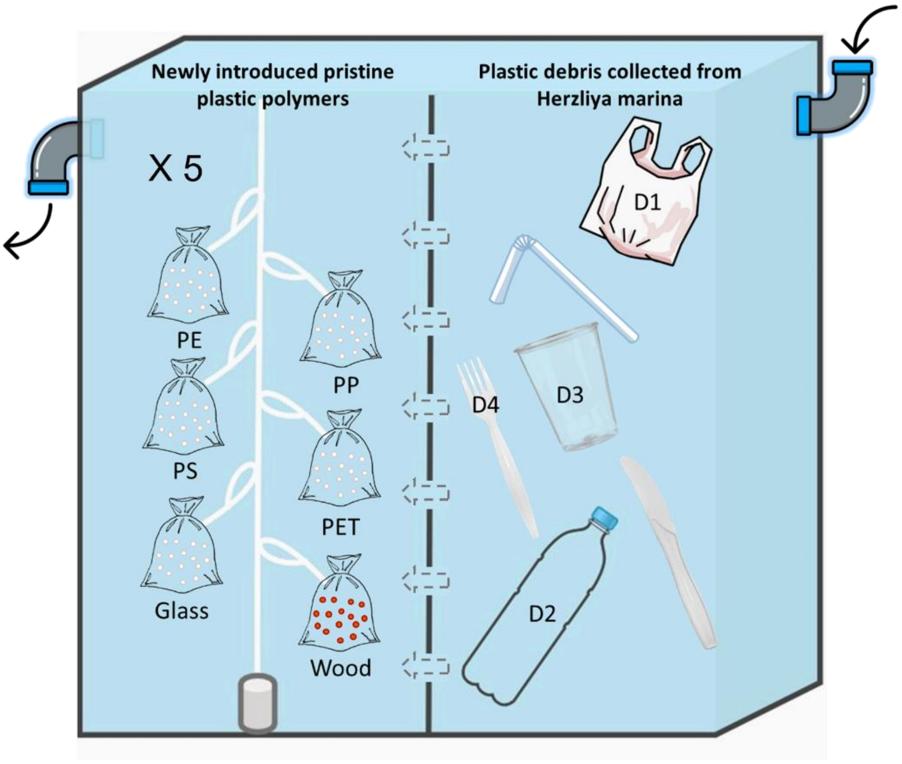


**Figure S1. Experiment setup.** The two aquarium compartments were separated by a polycarbonate barrier with round holes, 2 cm in diameter, 7.5 cm apart, enabling water interchange (dashed line). On the left compartment – organza mesh bags containing beads made of different plastic polymers, wood, and glass (5 replicates of each of 6 materials). On the right compartment – plastic items collected from the environment (represented by D1-D4 according to the items that were sampled). Water inflow is to the aquarium far right corner; water outflow is from the far-left corner. The second experiment included only PE, PET, glass, and wood on the left compartment.

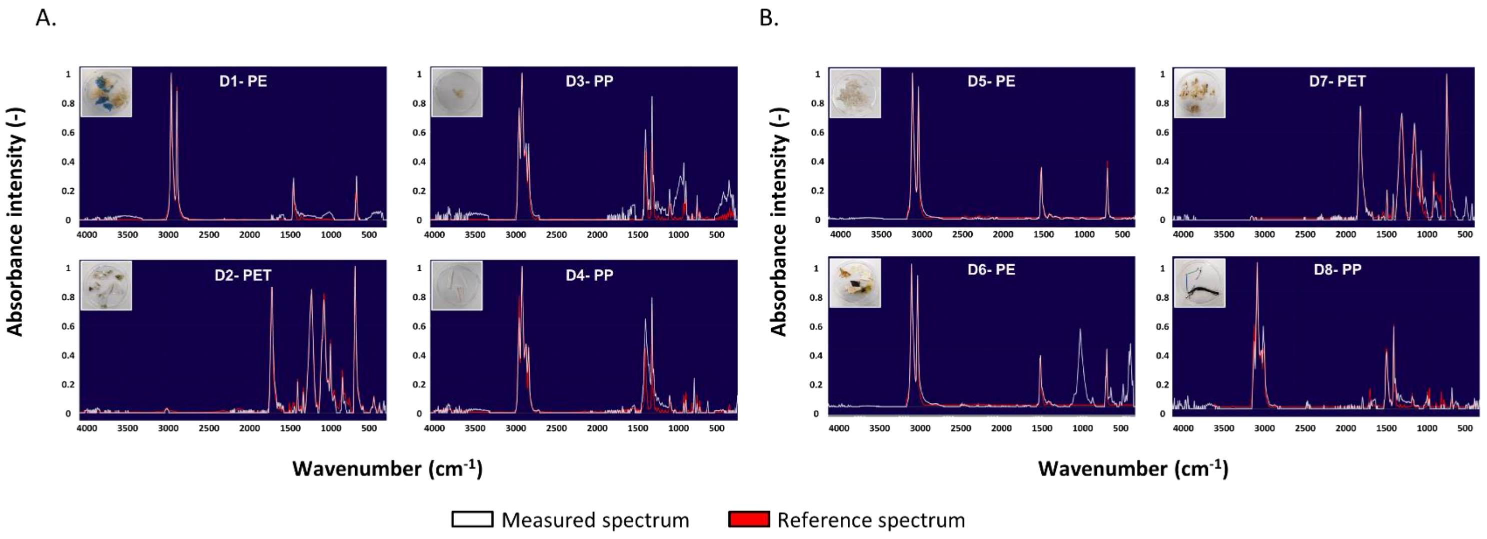


**Fig S2. FTIR spectral analysis of plastic debris.** A. Plastic debris items for the main experiment (D1-D4). B. Plastic debris items for the second experiment (D5-D8). The measured spectra are in white and the reference spectra are in red.

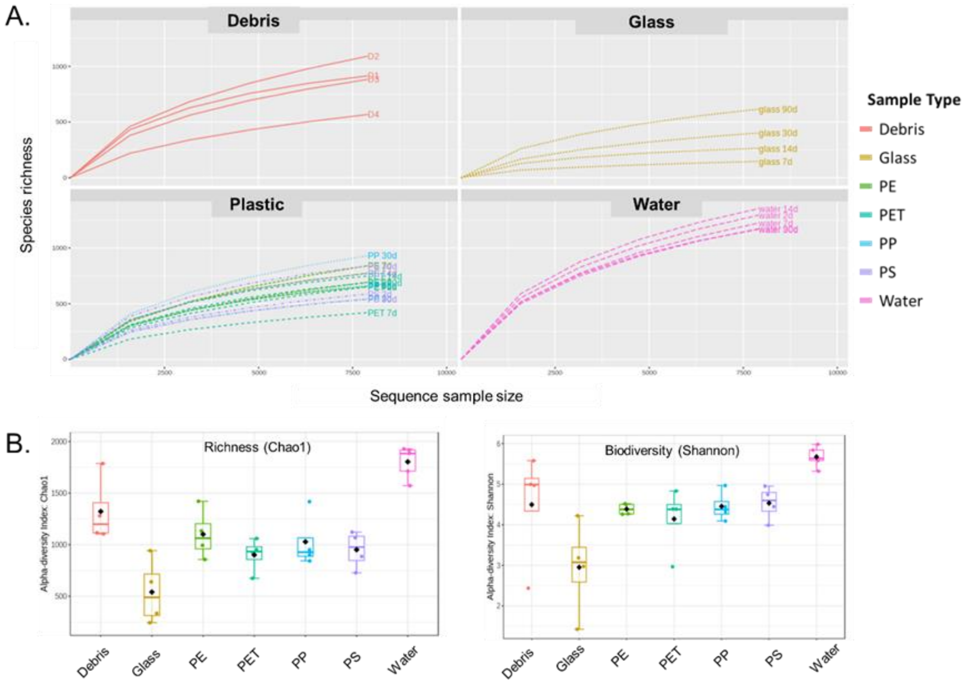


**Fig S3. Biodiversity rarefication curves and detailed biodiversity indexes.** A. Rarefication curves for Debris, Glass, Plastic and Water samples – species richness by sample size. B. Alpha Diversity (CHo1 and Shannon indexes) across all sample groups (Debris, Glass, PE, PP, PET, PS, and Water).

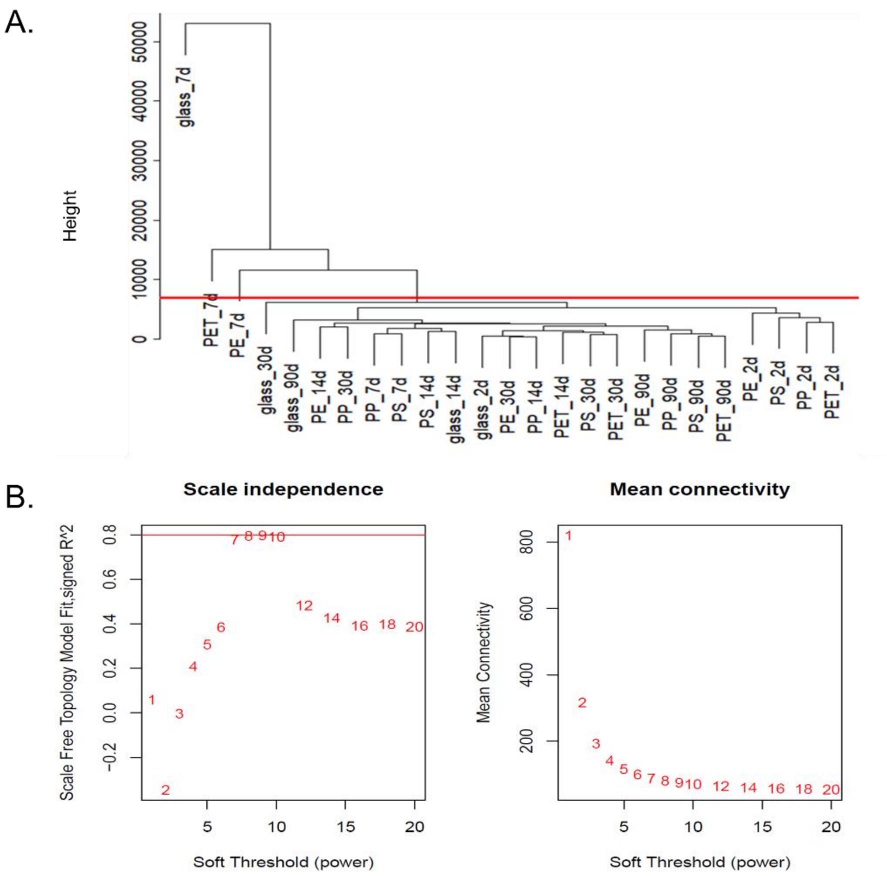


**Fig S4. Data pre-processing for WGCNA**. A. Sample dendrogram and trait heatmap after removing the outlying samples. The dendrogram illustrates the hierarchical clustering of samples. Glass, PET, and PE 7 days were removed from the analysis B. Scale Independence plot presents the scale-free topology fit index as a function of the soft-thresholding power. The red horizontal line represents the cutoff (R^2 = 0.80) for optimal scale independence. C. The mean connectivity plot presents the mean connectivity of the network as a function of the soft-thresholding power. The red labels indicate the corresponding powers used for analysis.

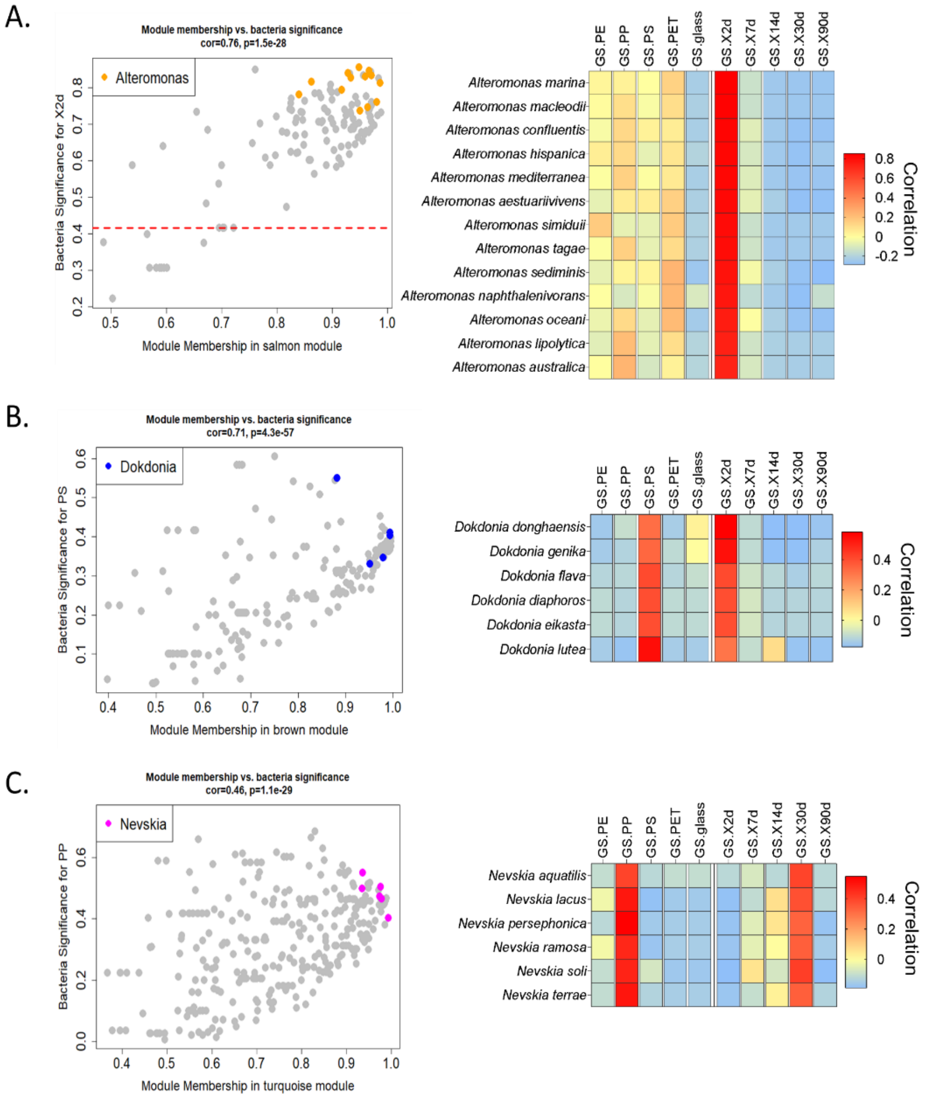


**Fig S5. Examples of module key genera.** Module membership values of the key genera – *Alteromonas* (A), *Dokdonia* (B), and *Nevskia* (C) are represented in the context of their associate modules: Salmon, Brown, and Turquoise accordingly (left side). The corresponding gene significance (GS) charts depicting the level of correlation of the key genera to all traits (surface type and time point) are presented on the right side. Dashed line indicates significance threshold (*p = 0.05*) to trait**.**

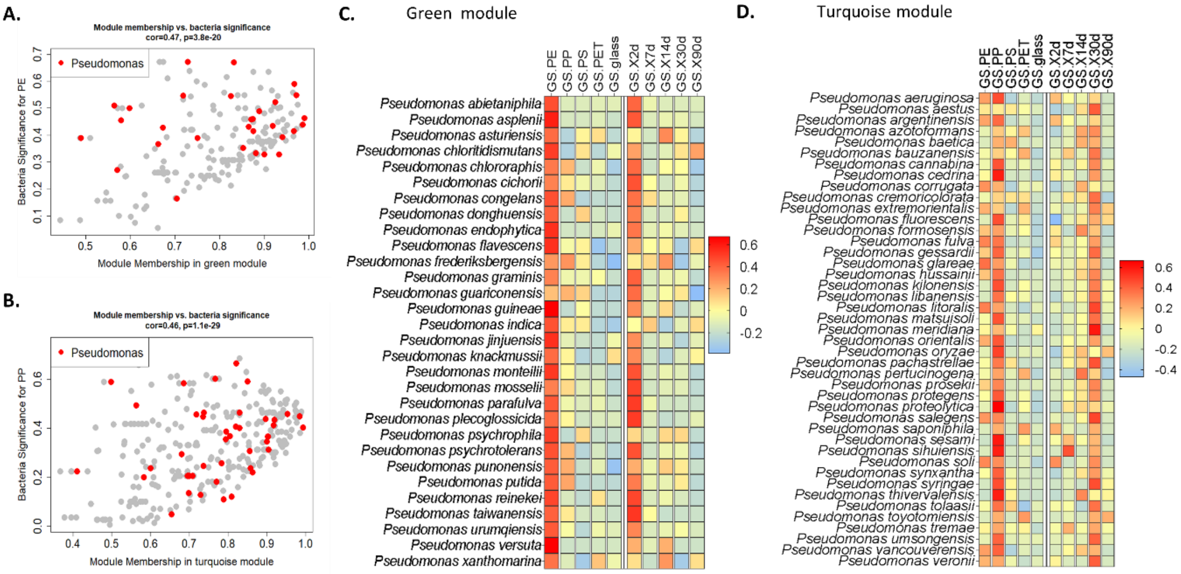


**Fig. S6. Pseudomonas as an example of a genus with succession sub-groups.** A, B. distribution of mapped species in modules Green (A) and Turquoise (B) by module membership and significance to the relevant trait (according to module eigengenes). Pseudomonas species are emphasized in Red. C, D. The corresponding gene significance (GS) heat maps for Pseudomonas, depicting the level of correlation to each trait (surface types and time points).

### Table S1. General run parameters

Experiment 1

| Sample | | Total reads | Mapped reads (LCA==0) | mapped reads (%)* | No. of species | RUN |
|---|---|---|---|---|---|---|
| 2 days | PE | 72,968 | 25,122 | 34.43 | 710 | 1 |
| | PP | 90,944 | 31,581 | 34.73 | 601 | 3 |
| | PS | 44,689 | 23,086 | 51.66 | 587 | 1 |
| | PET | 76,430 | 26,626 | 34.84 | 702 | 1 |
| | Glass | 2,277 | 1,337 | 58.72 | 79 | 3 |
| | Wood | 518 | 305 | 58.88 | 27 | 1 |
| | Water | 87,178 | 22,406 | 25.70 | 1,329 | 1 |
| 7 days | PE | 157,897 | 43,933 | 27.82 | 1,157 | 1 |
| | PP | 20,736 | 8,294 | 40.00 | 391 | 1 |
| | PS | 13,319 | 7,336 | 55.08 | 299 | 1 |
| | PET | 51,809 | 35,664 | 68.84 | 530 | 3 |
| | Glass | 78,470 | 73,857 | 94.12 | 245 | 1 |
| | Wood | 7,565 | 5,001 | 66.11 | 145 | 1 |
| | Water | 61,875 | 18,657 | 30.15 | 1,126 | 1 |
| 14 days | PE | 42,917 | 9,425 | 21.96 | 503 | 2 |
| | PP | 16,711 | 3,395 | 20.32 | 331 | 2 |
| | PS | 46,572 | 13,037 | 27.99 | 591 | 3 |
| | PET | 33,924 | 9,006 | 26.55 | 487 | 3 |
| | Glass | 11,132 | 8,487 | 76.24 | 167 | 3 |
| | Wood | 184 | 79 | 42.93 | 16 | 3 |
| | Water | 60,091 | 13,667 | 22.74 | 1,105 | 2 |
| 30 days | PE | 11,923 | 2,119 | 17.77 | 195 | 2 |
| | PP | 73,689 | 1,4833 | 20.13 | 773 | 2 |
| | PS | 57,996 | 11,460 | 19.76 | 613 | 3 |
| | PET | 28,734 | 5,374 | 18.70 | 260 | 3 |
| | Glass | 41,877 | 20,539 | 49.05 | 369 | 3 |
| | Wood | 25,350 | 9,455 | 37.30 | 309 | 3 |
| | Water | 38,608 | 10,380 | 26.89 | 818 | 2 |
| 90 days | PE | 62,930 | 9,483 | 15.07 | 436 | 3 |
| | PP | 42,039 | 10,364 | 24.65 | 437 | 3 |
| | PS | 43,490 | 9,751 | 22.42 | 340 | 3 |
| | PET | 60,616 | 12,099 | 19.96 | 482 | 3 |
| | Glass | 43,357 | 17,441 | 40.23 | 555 | 3 |
| | Wood | 11,253 | 2,662 | 23.66 | 204 | 3 |
| | Water | 64,132 | 18,742 | 29.22 | 1,138 | 3 |
| Debris | D1 | 29,231 | 7,940 | 27.16 | 556 | 1 |
| | D2 | 265,893 | 65,849 | 24.77 | 1,963 | 1 |
| | D3 | 71,037 | 17,864 | 25.15 | 761 | 1 |
| | D4 | 209,368 | 119,008 | 56.84 | 1,277 | 1 |

Experiment 2

| Sample | | Total reads | Mapped reads (LCA==0) | mapped reads (%)* | No. of species | RUN |
|---|---|---|---|---|---|---|
| 7 days | PE | 189,159 | 48159 | 25.46 | 1,133 | 4 |
| | PET | 3,880 | 1656 | 42.68 | 150 | |
| | Glass | 910 | 586 | 64.40 | 48 | |
| | Wood | 407,636 | 287567 | 70.55 | 1,078 | |
| | Water | 44,870 | 14261 | 31.78 | 836 | |
| 14 days | PE | 194,577 | 37580 | 19.31 | 901 | |
| | PET | 52,028 | 12728 | 24.46 | 746 | |
| | Glass | 688 | 238 | 34.59 | 28 | |
| | Wood | 74,247 | 52727 | 71.02 | 492 | |
| | Water | 14,2815 | 36235 | 25.37 | 1,666 | |
| 30 days | PE | 183,080 | 30897 | 16.88 | 1,189 | |
| | PET | 3,201 | 597 | 18.65 | 102 | |
| | Glass | 1,180 | 361 | 30.59 | 47 | |
| | Wood | 195 | 80 | 41.03 | 14 | |
| | Water | 120,956 | 25272 | 20.89 | 1,441 | |
| Debris | D1 | 3,833 | 706 | 18.42 | 69 | |
| | D2 | 1,7115 | 4166 | 24.34 | 312 | |
| | D3 | 2,112 | 431 | 20.41 | 69 | |
| | D4 | 762,64 | 18296 | 23.99 | 984 | |

Sequencing coverage averages (in reads), not including debris

Experiment 1
Average – 45,263 reads
STDV- 32,465

Experiment 2
Average- 94,628
STDV – 114,734
